# Network architecture of transcriptomic stress responses in zebrafish embryos

**DOI:** 10.1101/2024.06.30.601387

**Authors:** Kaylee Beine, Lauric Feugere, Alexander P. Turner, Katharina C. Wollenberg Valero

## Abstract

Protein-protein interaction (PPI) network topology can contribute to explain fundamental properties of genes, from expression levels to evolutionary constraints. Genes central to a network are more likely to be both conserved and highly expressed, whereas genes that are able to evolve in response to selective pressures but expressed at lower levels are located on the periphery of the network. The stress response is likewise thought to be conserved, however, experimental evidence for these patterns is limited. We examined whether the transcriptomic response to two environmental stressors (heat, UV, and their combination) is related to PPI architecture in zebrafish (*Danio rerio)* embryos. We show that stress response genes are situated more centrally in the PPI network. The transcriptomic response to heat was located in both central and peripheral positions, whereas UV response occupied central to intermediate positions. Across treatments, differentially expressed genes in different parts of the network affected identical phenotypes. Our results indicate that the zebrafish stress response has mostly conserved but also some stressor-specific aspects. These properties can aid in better understanding the organismal response to diverse and co-occurring stressors. Network position was further linked to the magnitude of fold changes of genes and types and number of linked phenotype components. Given the speed of contemporary changes in aquatic ecosystems, our approach can aid in identifying novel key regulators of the systemic response to specific stressors.

## Introduction

An organism’s capacity to perceive and adjust to environmental challenges plays a crucial role in upholding functional cellular homeostasis (López-Maury, Marguerat, and Bähler 2008). It also may align to the potential of species to respond to climate change by either acclimatizing to stressful conditions or by shifting their geographical distribution to more favorable habitats (Bernatchez et al. 2024). In recent years, there has been an increased focus on ectothermic organisms, which are heavily influenced by temperature dynamics (Rebolledo, Sgrò, and Monro 2021; Sokolova 2021). In addition to the increased incidences of abnormally high temperatures during heat waves, organisms must also deal with stress induced by other aspects of the abiotic environment, such as UV radiation (UVR) experienced from sunlight (Dahms and Lee 2010), which is also expected to become more intense due to the effect of greenhouse gases (Eleftheratos et al., 2022). However, aquatic organisms may show more complex responses to combined stressors due to interactive effects (Côté, Darling, and Brown 2016; Piggott, Townsend, and Matthaei 2015). For instance, UVR and heat can synergistically or antagonistically impact fish biology (Cramp et al. 2014; Alves and Agustí 2020; Icoglu Aksakal and Ciltas 2018; Feugere et al., n.d.). UVR has been shown to affect biological processes in ectotherms from fish (Ripley et al. 2023), to insects (Jørgensen et al. 2022), to amphibians (Londero, Santos, and Schuch 2019), including growth, homeostasis, DNA damage and repair. In some cases, this results in negative consequences for the performance and fitness of the organisms. For example, heat has been shown to strongly affect the integrity of cellular structures (Porcelli et al. 2015), denature DNA repair proteins (Dahms and Lee 2010), and increase ROS production (Mariana and Badr 2019) in fish. Similarly, UVR is known to cause damage to proteins, lipids, and DNA in fish (Griffiths et al. 1998; Dahms and Lee 2010; Alves and Agustí 2020). Fish embryos are more vulnerable to heat and UVR than their adult counterparts, as developmental functions are impacted by mutations and heat stress (Alves and Agustí 2020; Ripley et al. 2023). Heat stress combined with UVR during early-life development can increase malformations and mortality and so reduce the fitness of fish, which lay eggs in shallow, often warm, UVR-exposed waters (Yabu, Todoriki, and Yamashita 2001; Dahms and Lee 2010; Alves and Agustí 2020; Lundsgaard et al. 2020; Feugere et al., n.d.). Both ultraviolet A (UVA, 315-400 nm) and B (UVB, 280-315 nm) cause damage to macromolecules (Alves and Agustí 2020; Dahms and Lee 2010; Rastogi et al. 2010), but UVA (and blue light) can activate repair mechanisms to reverse DNA lesions (Banaś et al. 2020; Dong et al. 2007; Rastogi et al. 2010). As such, UVA following UVB exposure can partially rescue malformed zebrafish embryos, confirming the photorepair capacity of this species (Dong et al. 2007; Feugere et al., n.d.). In addition, sublethal heat stress during development can have a hormetic effect, protecting zebrafish embryos grown under high temperatures against further damage through UVR (Feugere et al., n.d.). In summary, there is precedence for using UVR combined with heat exposure during zebrafish development as a template to study the effects of different environmental stressors on zebrafish.

One central role in cellular acclimation to short- or long-term environmental stressors lies in the modulation of gene expression, where extensive regulation takes place at both the transcriptional and post-transcriptional levels (López-Maury, Marguerat, and Bähler 2008). The control of gene expression in response to a stressor is both tightly regulated and reversible, depending on the type of stress and the organism (de Nadal, Ammerer, and Posas 2011). For instance, warm temperatures during embryonic development caused later-life effects on the transcriptomic network organization in laboratory lines of multiple fish species, including zebrafish, increasing its network entropy (Scott and Johnston 2012; Ripley et al. 2023). The stress response is regarded to be adaptive, and depending on the organism and the type of stress, can either be generically shared by multiple stressors or be specific to a particular stressor (de Nadal, Ammerer, and Posas 2011; Wollenberg Valero et al. 2022). Any stressor-specific response may trigger deleterious trade-offs within multiple stressor environments, which is why the general stress response is thought to have an evolutionary conserved component (de Nadal, Ammerer, and Posas 2011; Burton et al. 2021).

Genetic networks, such as protein-protein interaction (PPI) networks, describe functional interactions of proteins within the cell. Genetic networks are often used in human research (Rual et al. 2005; Luck et al. 2020). Additional studies have focused on network evolution in fungi (Joy et al. 2005; Usaj et al. 2017; Lorena Ament-Velásquez et al. 2022; Wollenberg Valero 2020) and bacteria (Philippe et al. 2007; Crombach and Hogeweg 2008), with very few studies pertaining to vertebrate animals (but see (Wollenberg Valero et al. 2014, 2022) for examples of climate adaptation-related networks). However, a few general patterns are emerging; such as that of genes towards the center of the network being most highly conserved, and those positioned intermediately having the highest number of connections with other nodes, whereas both constraint and pleiotropy are lowest at the network periphery (also see (Wollenberg Valero 2024). While these properties pertain to adaptive processes accumulated over multiple generations, they nonetheless also affect gene expression, as functional connections in a PPI denote real-time interactions between gene products and can thus link evolution to function. Networks can explain such emergent properties which are not apparent when studying genes or pathways individually (Bhalla and Iyengar 1999). Highly-expressed genes are under strong natural selection for correct folding and function, and thus evolve slowly, a phenomenon known as the “E-R anti-correlation” (Joy et al. 2005). These genes, located centrally in the interactome, are functionally constrained as they are essential for cell survival (Alvarez-Ponce et al. 2016; Wollenberg Valero 2020; Jeong et al. 2001), and show a lower evolutionary rate (Alvarez-Ponce, Feyertag, and Chakraborty 2017). Gene expression is lower in highly pleiotropic genes, a phenomenon dubbed the “cost of complexity” (Fisher, 1930; Orr, 2000). The number of protein-protein interactions is lower in proteins whose evolutionary rate is higher (Alvarez-Ponce, Aguadé, and Rozas 2009; Alvarez-Ponce and Fares 2012). In vertebrates, a set of ∼1000 genes identified to adapt to climatic gradients is to a large part differentially up- or downregulated (38%, and 22.5%, respectively) in response to short-term environmental stressors such as heat or cold stress (Wollenberg Valero et al. 2022). Of these genes, 25% are also “wider definition” housekeeping genes with intermediate levels of expression (Eisenberg and Levanon 2013; Wollenberg Valero et al. 2022). It is therefore of interest to study the location of genes involved in the stress response within PPI networks, particularly with respect to combined stressors, in order to better understand the interplay between expression levels, stress response, and network position.

In this paper, we analyze a transcriptomic dataset previously generated from developing zebrafish embryos (Feugere et al., n.d.) exposed to heat stress (TS) and UV radiation (UV). Our analysis includes three treatment comparisons: one for each stressor against a control, and one comparing UV following TS to UV exposure alone.

Four specific hypotheses are tested. **Hypothesis (i)** is that genes linked to the stress response in terms of their Gene Ontology (GO), cluster centrally within the zebrafish PPI network, reflecting both a high degree of evolutionary constraint and the potential for high gene expression expected under such a central, conserved stress response. **Hypothesis (ii)** is that this pattern also applies to genes differentially expressed in response to two distinct stressors (TS and UV). Under **hypothesis (iii),** we expect that some genes are expressed only when TS follows UV (synergistically), and others are only expressed under each single stress condition (antagonistically), expected to be located in peripheral regions in the network. **Hypothesis (iv)** We anticipate that phenotypic outcomes (such as gene ontologies, and genes’ links to phenotypes and diseases), along with the relative changes in gene expression, are influenced by their positions within the Protein-Protein Interaction (PPI) network. Genes located in hub or intermediate positions of the PPI are expected to show both higher expression levels and evolutionary constraint, and due to pleiotropy may have significant phenotypic effects. In contrast, peripheral genes, which generally have lower levels of expression and constraint, may exhibit more variable phenotypic effects and larger fluctuations in expression changes.

## Materials and Methods

All transcriptomic and phenotypic data used in this study were obtained from (Feugere et al., n.d., 2023), but reanalyzed to make novel comparisons. A more thorough description of methods used such as animal husbandry and setup of the UVR treatment comparisons can be found therein. All experiments were approved by the Ethics committee of the University of Hull (FEC_2019_194 Amendment 1). Briefly, zebrafish embryos (2 to 3.3 hours post fertilization, hpf) were exposed for 24 h to either constant control temperature alone (27°C), or to treatment comparisons consisting of thermal stress (19 peaks of sublethal 32°C), 24 hof constant temperature followed by UV radiation damage/repair assay (Dong et al. 2007), or both (first 24 h of heat peaks followed by UV radiation damage/repair assay). The UV assay consisted of 6 min of UVB followed by 15 min of UVA. Embryos were subsequently humanely sacrificed by snap-freezing at -80°C, followed by RNA extraction, cDNA synthesis, and Illumina sequencing (Feugere et al. 2023). Differential gene expression was performed with DESeq2 v1.28.1 (Love, Huber, and Anders 2014) following read quality control using *fastp* v0.23.1 (Chen et al. 2018) and mapping against the zebrafish reference genome with the STAR v2.6.1 aligner (Dobin et al. 2013). To identify differentially expressed genes (DEGs), we compared various treatment comparisons to a control. We abbreviated each comparison for clarity: “**TS**” represents the comparison between the thermal stress treatment and the control, “**UV**” denotes the UVB/UVA treatment to the control, and “**TS->UV**” refers to the comparison of thermal stress followed by UVB/UVA damage/repair assay against UVB/UVA damage/repair assay without prior thermal stress. DEGs were determined to be significant when p-adj < 0.05 and absolute fold-change |FC| ≥ 1.5. This paper, compared to (Feugere et al., n.d., 2023), outlines gene expression differences as a result of thermal stress and UV exposure, and specifically characterizes network statistics using experimental data.

The position of a node in a network is dependent on the overall structure of the network, since statistical properties of experimentally-determined nodes are computed relatively to all other nodes in a network. In this study, two reference networks are used: **(i)** The first reference network, a complete PPI network for zebrafish, was a filtered and re-annotated STRING network obtained from Fernando and colleagues (Fernando, Mabee, and Zeng 2020) which contained 14,677 genes (nodes) and 247,439 interactions (edges), with an edge evidence score of 0.9 out of 1, hereafter referred to as the zebrafish **interactome**. This PPI was lacking 1006 genes (nodes) from our experimentally expressed genes. Of these, 873 genes and their interactions (edges) were manually imported through STRING (https://string-db.org/). The missing genes were fitted into the network through matching the connecting genes found in the original Fernando and colleagues network via .sif format. **(ii)** Since our experimental data was obtained from a transcriptomics experiment, only genes that are expressed in the developing embryo around 24 hpf would be likely to perform functional interactions with one another and may therefore constitute another appropriate reference network. The second reference network was therefore the final interactome, but filtered by all genes which were not expressed in any of our experiments with a read count of >10. This is here referred to as the **expressed network**. This network contained 10,953 nodes and 159,302 edges. For each reference network, any other single, unconnected nodes were removed from the analysis as their network properties would not be known.

Within any PPI network, nodes represent proteins, and edges connect interacting proteins. We previously demonstrated that the topology of nodes within a network can be characterized by decomposing a node’s position into three statistical parameters: Neighborhood Connectivity (**NC**, with the highest values characterizing nodes located intermediately in the network, and with the highest number of edges to other nodes), Average Shortest Path Length (**ASPL**, with highest values characterizing nodes peripherally located in the network and with few connections to other nodes), and Betweenness Centrality (**BC**, with highest values characterizing nodes central to the network) (Wollenberg Valero, 2020). These three network statistical parameters were calculated for all nodes in the reference networks and all DEGs from **TS**, **UV**, and **TS->UV** overlaid over these, using the Network Analyzer function in Cytoscape (v.3.10.0, (Shannon et al. 2003; Wollenberg Valero 2020)). These three metrics were then used to bin each node into one of three categories using Discriminant Function Analysis (DFA). All genes in the network and DEGs were assigned to peripheral nodes (P), intermediate nodes (I) and hub nodes (H), by considering their individual ASPL, NC and BC values, respectively, as previously described in Wollenberg Valero (2020) and Wollenberg Valero (2024).

Not only do network node statistical parameters change based on the size and content of the reference network, but also the adjusted p-value of differential gene expression differs based on the number of comparisons performed (genes in a dataset). Therefore, for each comparison between an experimental condition and a reference network, the p-value for differential gene expression was adjusted based on the size of the reference network (either the interactome; or the expressed network), using the *fdr* function (RStudio v.3.4.0, (R Core Team 2024)) to control for false positives.

Subsequently, DEGs in each of the experimental comparisons TS, UV, and TS->UV were overlaid over each of the two reference networks in Cytoscape. The following set of comparisons were made: **(i)** between the reference interactome and the stress GO term reference network; **(ii)** between the reference interactome and DEGs from the experimental data; **(iii)** the expressed network and the significant experimental DEGs. The data were not normally distributed (all comparisons p<0.001) according to Kolmogorov-Smirnov tests. Statistical tests were performed with SPSS v.27 (IBM Corp. 2020).

To test for **Hypothesis 1**, the zebrafish interactome was compared to the stress GO term network, using non-parametric Kruskal-Wallis tests followed by pairwise Mann-Whitney Wilcoxon tests followed by pairwise Mann-Whitney Wilcoxon tests between network statistical parameters of DEGs vs. the interactome or the expressed network, adjusted by Bonferroni correction. Due to the high number of genes, we calculated the partial ή^2^ for Cohen’s h statistic effect size (Kassambara 2023). The effect size was small if it was < 0.06; moderate if between 0.06 and 0.14; and large if ≥ 0.14.

To test for **Hypothesis 2**, we tested whether the transcriptomic responses to each treatment comparison in developing zebrafish are constrained to specific parts of the PPI using Kruskal-Wallis tests.

We then tested for **Hypothesis 3**, by selecting synergistic (combined treatment) and antagonistic DEGs (only in single stressor comparisons) as shown in Fig. 2 and comparing their network statistical properties to the interactome and expressed network, respectively as described for Hypothesis 1 and 2.

To test for **Hypothesis 4**, the significant DEGs per treatment comparison (TS, UV, TS->UV) were assessed for biological pathways, phenotype and diseases using the ShinyGO (v.8.0) features ‘*GO biological processes*’, ‘*ZFIN phenotype*’ and ‘*ZFIN diseases*’ respectively (Ge, Jung, and Yao 2020) and compared based on their location within the zebrafish interactome. DEG features shared by treatment comparisons were visualized using chord diagrams with single DEG features shown in supplementary data. Patterns of association between node categories or treatment comparisons were identified using chord diagrams and ShinyGO data.

All graphs were prepared in R (v.4.3.0) using *vennDiagram* (Chen 2022), *plotly* (Sievert 2020) for scatter plots, *ggplot2* (Wickham et al. 2020) for error plots, and *circlize (Gu et al. 2014)* for chord diagrams.

*ZFIN* phenotype and disease features can give insights on the normal functioning of genes, despite that the data is obtained from cases where the gene is malfunctioning or mutated (Sprague et al. 2001; Bradford et al. 2022). In order to integrate these ShinyGO features with experimentally-measured phenotypic outcomes in zebrafish embryos responding to the three treatment comparisons, data on sum of malformations and shortest embryo length (measured as a straight line from head to tail tip) were collected from 4 days post fertilisation (dpf) larvae (Feugere et al., preprint and additional unpublished data). Malformation scores were the sum of the presence or absence of three malformation types: bent spine, tail malformation, and pericardial edema. Shortest embryo length was the shortest distance from the head to the tail of the embryo measured with NIH ImageJ (Schneider, Rasband and Eliceiri, 2012). These phenotypic responses were, together with NC, ASPL and BC, standardized from -1 to 1 to be able to compare their relative changes across treatment comparisons.

## Results

### Hypothesis (i): Stress response GO term genes are more central to the PPI network

Based on classification using BC, NC, and ASPL, the zebrafish interactome (Fig. 1A) is characterized by mostly peripheral (62.1%,), followed by intermediate (35.7%) and central (2.13%) nodes. The stress GO term genes shared these overall proportions, but were located with less peripheral (58.9%), but more intermediate (37%) and central (4.19%) node positions within the interactome (pie chart percentages found in Supplementary Table S6). Fig. 1B shows the pairwise comparison between network statistics of the interactome compared to the placement of the stress GO term genes within it. The GO term genes occupied a significantly lower ASPL (KW-H: 122.454, p<0.001, with small effect size), higher but not significantly different NC, and significantly higher BC than the interactome (KW-H: 127.626, p<0.001, with small effect size) (*Supplementary Table S1*).

**Fig. 1:**
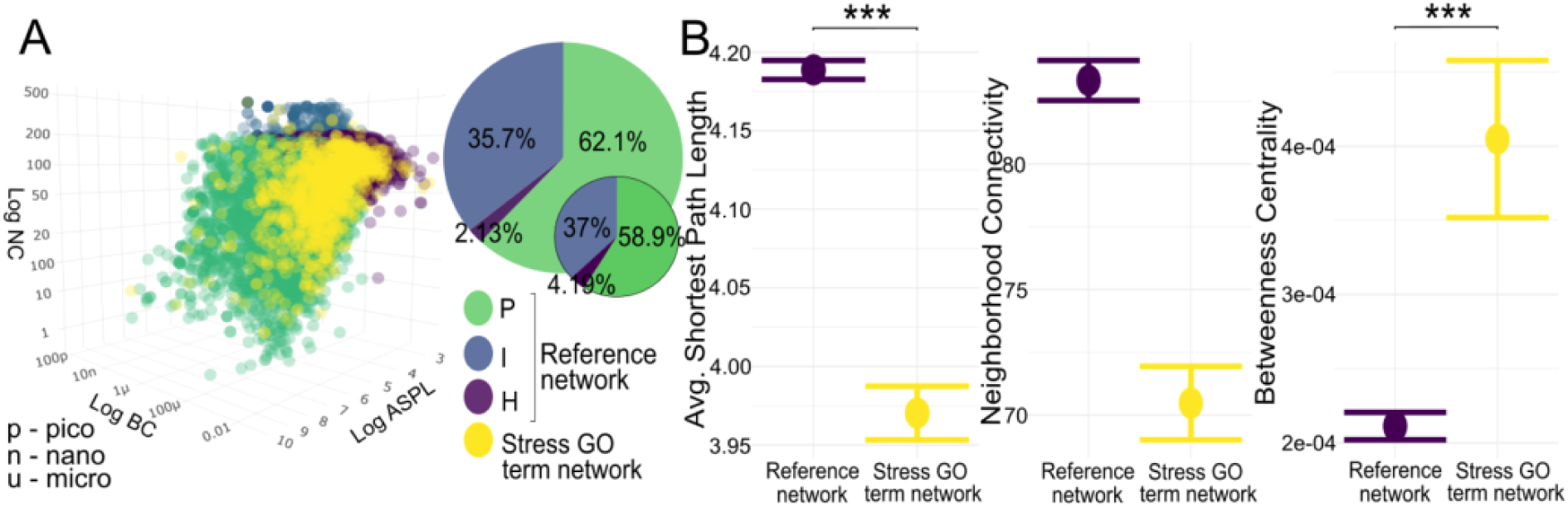
The position of stress GO term genes within the zebrafish interactome. (A) Distribution of interactome nodes within network parameter space represented by ASPL, NC, and BC, with node categories in green (P nodes), blue (I nodes), and purple (H nodes). Stress GO term genes are shown in yellow. Inset pie charts represent node category distribution (large: interactome; small: stress GO term genes). (B) Means with standard error plots of ASPL, BC and NC for the two gene sets. Significance levels of pairwise Mann-Whitney U tests are indicated with asterisks as p <0.0001: ***, p<0.001: **; p<0.05: *.

### Hypothesis (ii): Experimental stress response DEGs are more central to the interactome

DEGs from the three different treatment comparisons TS, UV and TS->UV were visualized and further subsetted based on a Venn diagram (Fig. 2). We highlighted genes shared between two or more stressors compared to the genes differentially expressed in only a single stressor, genes responding to all stressors, as well as antagonistic genes responding to both TS and UV independently but not in the combined treatment comparison TS->UV, and synergistic genes responding only in the TS->UV treatment comparison but not in individual treatment comparisons. Overall, 71 genes were shared between all three treatment comparisons (TS, UV, TS->UV). 187 genes were shared between UV and TS->UV, 121 genes were shared between TS and TS->UV, and 135 genes were shared antagonistically. In addition, 134 genes were only differentially expressed in TS but in no other combination, 1720 genes were only differentially expressed in UV but no other combination, and 287 were only differentially expressed in TS->UV synergistically, but not in the individual TS and UV treatment comparisons.

**Fig. 2:**
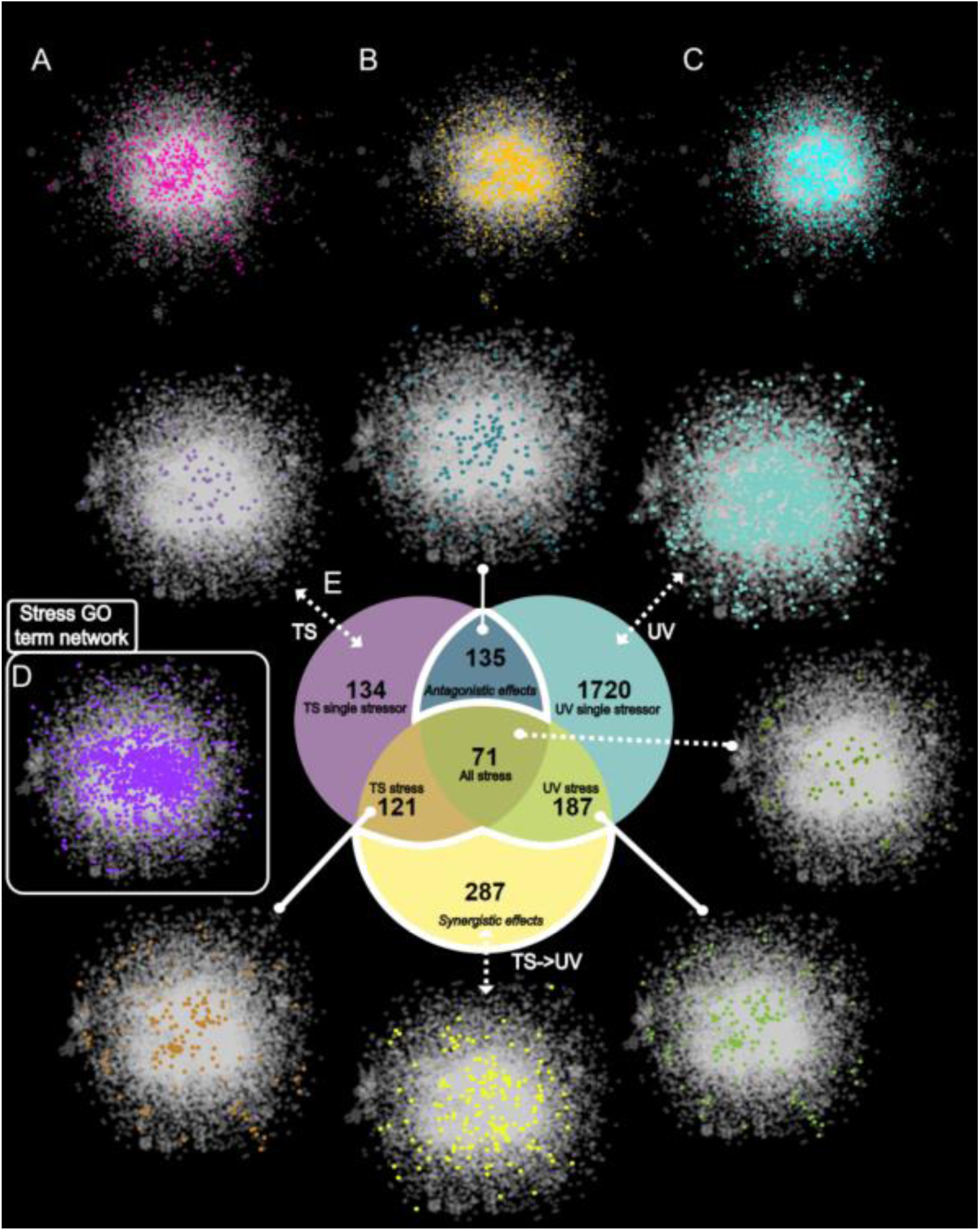
Visualization of significant DEGs in response to thermal stress, UV exposure, and UV exposure following thermal stress, highlighting differences (white) and commonalities (gray). The networks displayed in reference to the Venn diagram show the positioning of the genes relative to the zebrafish interactome and the stress GO term network. A - thermal stress genes, B - UV stress genes, C - Thermal stress followed by UV stress genes, D - Stress GO term network, E - Venn diagram with corresponding single stressors/combination stressors as isolated by overlapping genes. All genes (dark green), TS-responsive DEGs (brown), antagonistic DEGs (dark blue), TS-only DEGs (lilac), UV-only DEGs (light blue), and synergistic DEGs (yellow).

Overall, the relative position of all experimental DEGs across treatment comparisons TS, UV and TS->UV significantly differed from that of the zebrafish interactome (ASPL KW-H: 20.778, p<0.001; NC KW-H = 19.837, p<0.001; BC KW-H: 19.795, p<0.001, with small effect size for all predictors, Fig. 3, Table S2). Pairwise post hoc tests (Supplementary Tables S2.1 - S2.3) revealed that TS did not have a significantly different ASPL compared to the interactome, but had both a lower NC (W=630.619, Padj.=0.01) and higher BC (W=-525.45, Padj.=0.02).

**Fig. 3:**
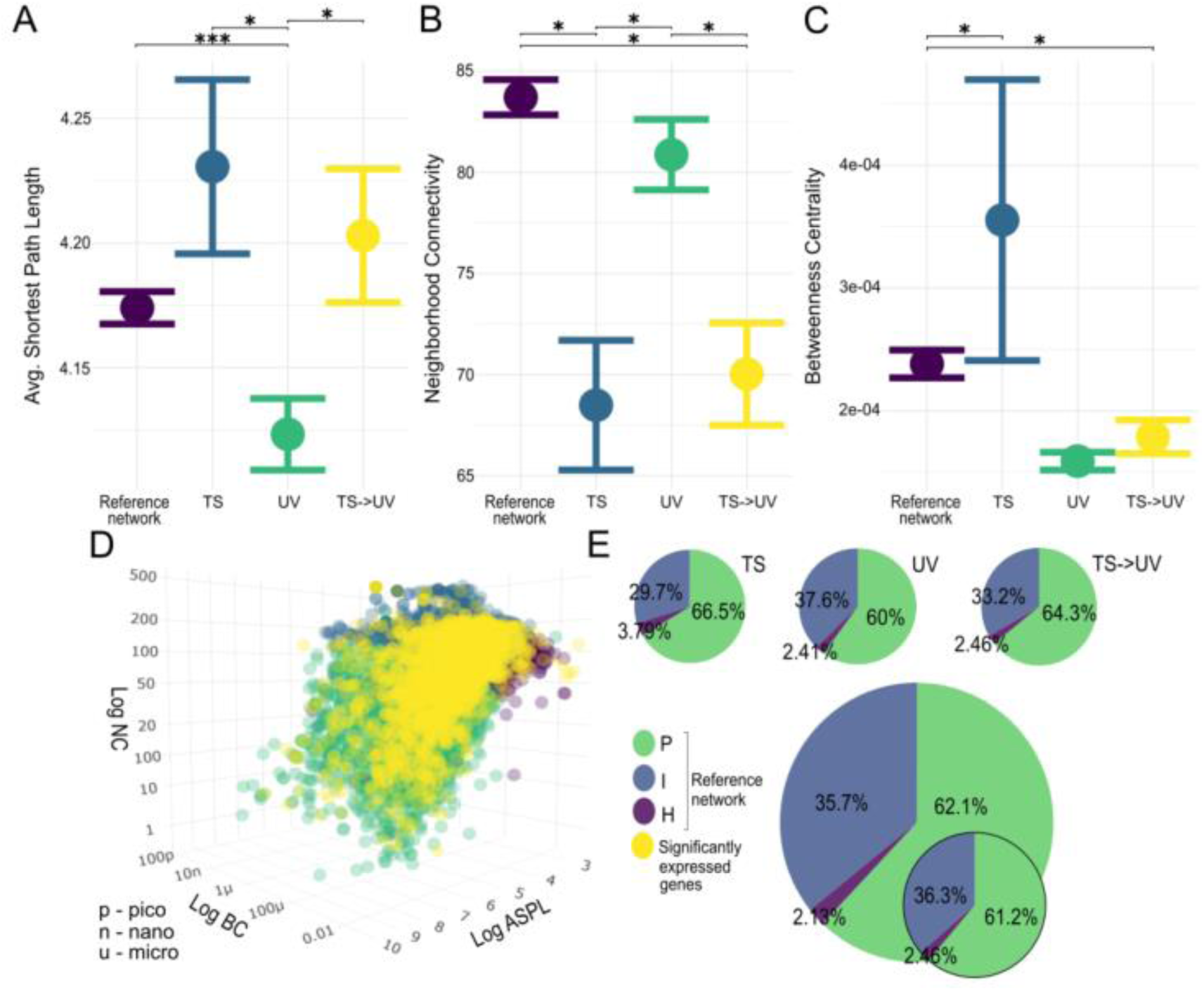
The position of experimental DEGs within the zebrafish interactome. (A-C) Means with standard error plots of ASPL, BC and NC for each gene set’s network dimensions with significance levels of pairwise Mann-Whitney U tests indicated with asterisks as p <0.0001: ***, p<0.001: **; p<0.05: *. The reference network is the interactome and TS, UV and TS->UV represent treatment comparison DEGs. (D) Distribution of interactome nodes within network parameter space represented by NC, ASPL, BC, with node categories shown in green (P nodes), blue (I nodes), and purple (H nodes). Experimental DEGs are shown in yellow. (E) Inset pie charts represent node category distribution (large: interactome; smaller: experimental DEGs, smallest: experimental DEGs per treatment comparison).

Correspondingly, it had 4.4%, (n=20 genes) more P nodes, 6%, (n=27) less I nodes, and 1.33%, (n=6) more H nodes compared to the interactome (Table S2.1 - S2.3). DEGs from the UV treatment comparison had a significantly lower ASPL (W=420.675, Padj.<0.001), and no significant difference in NC and BC. As expected, it had 2.1% (n=44) less P nodes, but also 1.9% (n=40) more I and 0.33% (n=6) more H nodes (pie chart percentages found in *supplementary table S6*). The combined TS->UV treatment comparison did not have a significantly different ASPL compared to the interactome, but a lower NC (W=453.921, Padj.=0.04), and higher BC (W=-481.9, Padj.=0.008). In accordance with this, the number of P nodes was 2.2% (n=15) higher, the number of I nodes 2.5% (n=17) lower and there were 0.33% (n=2) more H nodes (Table S2.1 - S2.3).

### Hypothesis (ii) The topology of DEGs shows the same pattern relative to the expressed portion of the network

The genes expressed during the experiment differed in their network topology from the interactome. The number of H nodes expressed in this part of the interactome was 1.88% (n=205 genes) lower, and most importantly, the number of P nodes was 36.4% (n=3987) lower in favor of 38.3% (n=4194) more I nodes than the zebrafish interactome. Network statistical parameters of experimental DEGs within this expressed network significantly differed between treatment comparisons in ASPL (KW-H: 13.226, p-value: 0.004) and NC (KW-H: 20.253, p-value: <0.001), but not BC (KW-H: 6.297, p-value: 0.098, with small effect size for all predictors, Fig. 4, Table S3). Consequently, no pairwise post hoc tests for BC were performed. In addition, while ASPL was a significant predictor in the overall analysis, treatment comparisons did not significantly differ in ASPL in post hoc tests (Supplementary Table S3.1). However, DEGs from the TS treatment comparison had significantly lower NC (W=504.88, Padj.=0.009) than the expressed network (Supplementary Table S3.1). Compared to the expressed reference network, only one more H node was activated (0.2%,), but I nodes decreased by 7.2% (n=33) in favor of 7.07% (n=33) more P nodes (Supplementary Table 5.1). DEGs from the UV treatment comparison did not differ in ASPL, NC or BC compared to the expressed network. However one more H node was activated (0.059%, n=1), and 1% (n=23) less P nodes were activated in favor of 1%, (n=21) more I nodes (Supplementary Table 6). DEGs from the TS->UV treatment comparison had significantly lower NC than the expressed network (W=362.267, Padj.=0.039; Fig. 4B; Table S3). There were only 2 less H nodes (-0.25%), but a decrease in I nodes of 3.3% (n=22) and a concomitant increase in P-nodes of 3.6% (n=24, Supplementary Table 6).

**Fig. 4:**
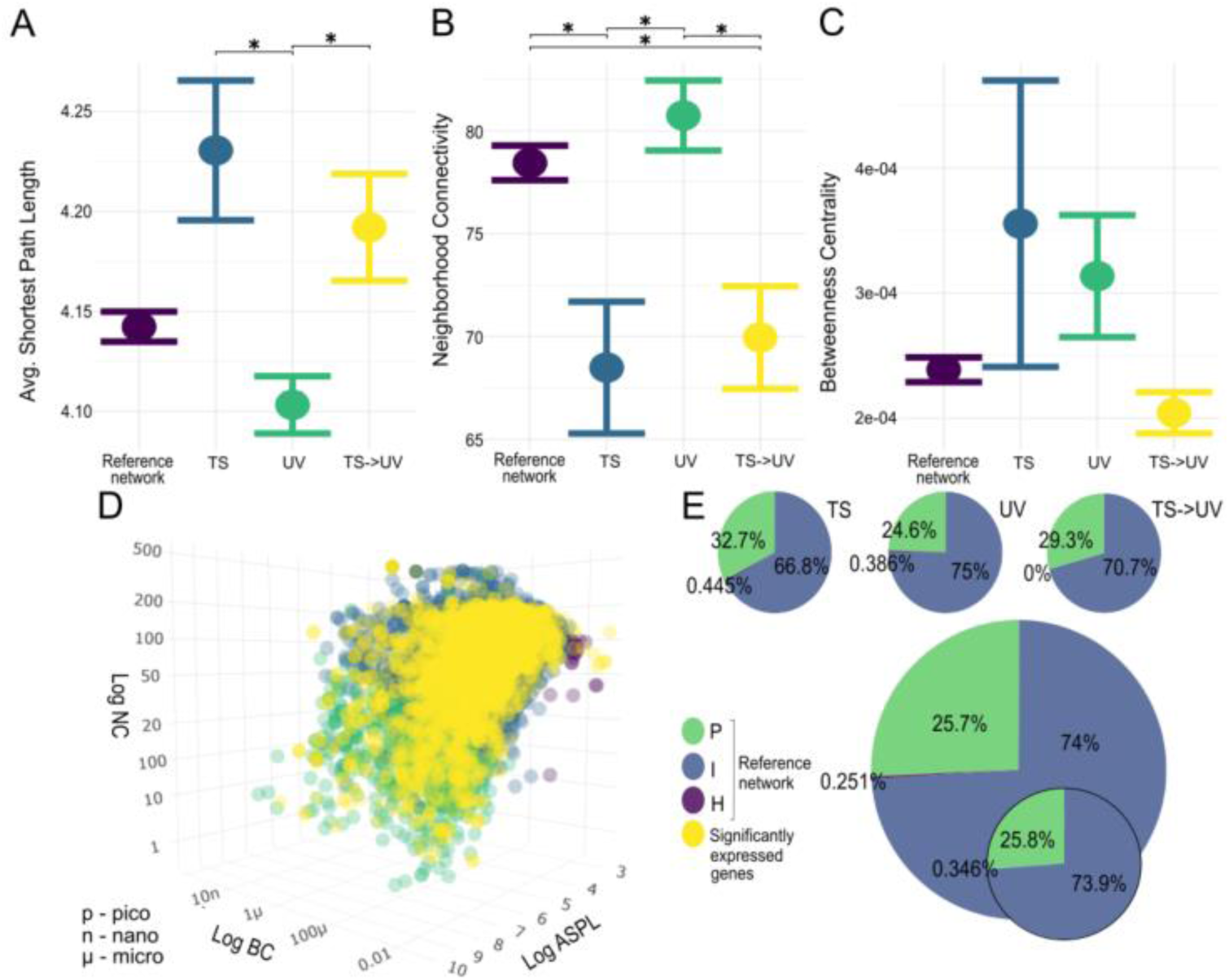
The position of experimental DEGs within the expressed network. (A-C) Means with standard error plots of ASPL, BC and NC for the compared gene sets’ network dimensions with significance levels of pairwise Mann-Whitney U tests indicated with asterisks as p <0.0001: ***, p<0.001: **; p<0.05: *. The reference network is constructed from genes expressed during the experiments and TS, UV and TS->UV represent treatment comparison DEGs. (D) Distribution of expressed network nodes within network parameter space represented by NC, ASPL, and BC, with node categories shown in green (P nodes), blue (I nodes), and purple (H nodes). Experimental DEGs are shown in yellow. (E) Inset pie charts represent node category distribution (large:expressed genes; smaller: experimental DEGs, smallest:experimental DEGs per treatment comparison).

### Hypothesis (iii): Combined stressors activate more central nodes while stressor-unique DEGs are more peripheral

Neither synergistic (only present in TS->UV) or antagonistic DEGs (in both TS or UV comparisons but not in TS->UV) differed by node statistics (Supplementary Fig S1, Supplementary Table S4). Nonetheless, the synergistic gene set had 3.7% (n=11) more P genes and 3.5% (n=10) less I genes compared to the interactome (Supplementary Table S6). The antagonistic gene set had 7.9% (n=11) less I nodes and 6% (n=8) more P nodes relative to the interactome (Supplementary Table S6). In addition, network statistical parameters did not differ significantly between DEGs in the TS treatment comparison and the interactome (Fig. 5, Supplementary Table S5), but TS had +0.92% (n=8) more P nodes and 6.7% (n=9) less I nodes (Supplementary Table S6). However, UV as a single stressor activated significantly more 2.7%, (n = 8) I nodes and significantly less P nodes 2.7%, (n=5) than the interactome, supported by both lower ASPL (H = 12.416, p <0.001, small effect size, Figure 5, Supplementary Table S5) and lower BC (H = 16.584, p <0.001, small effect size, Figure 5, Supplementary Table S5).

**Fig. 5.**
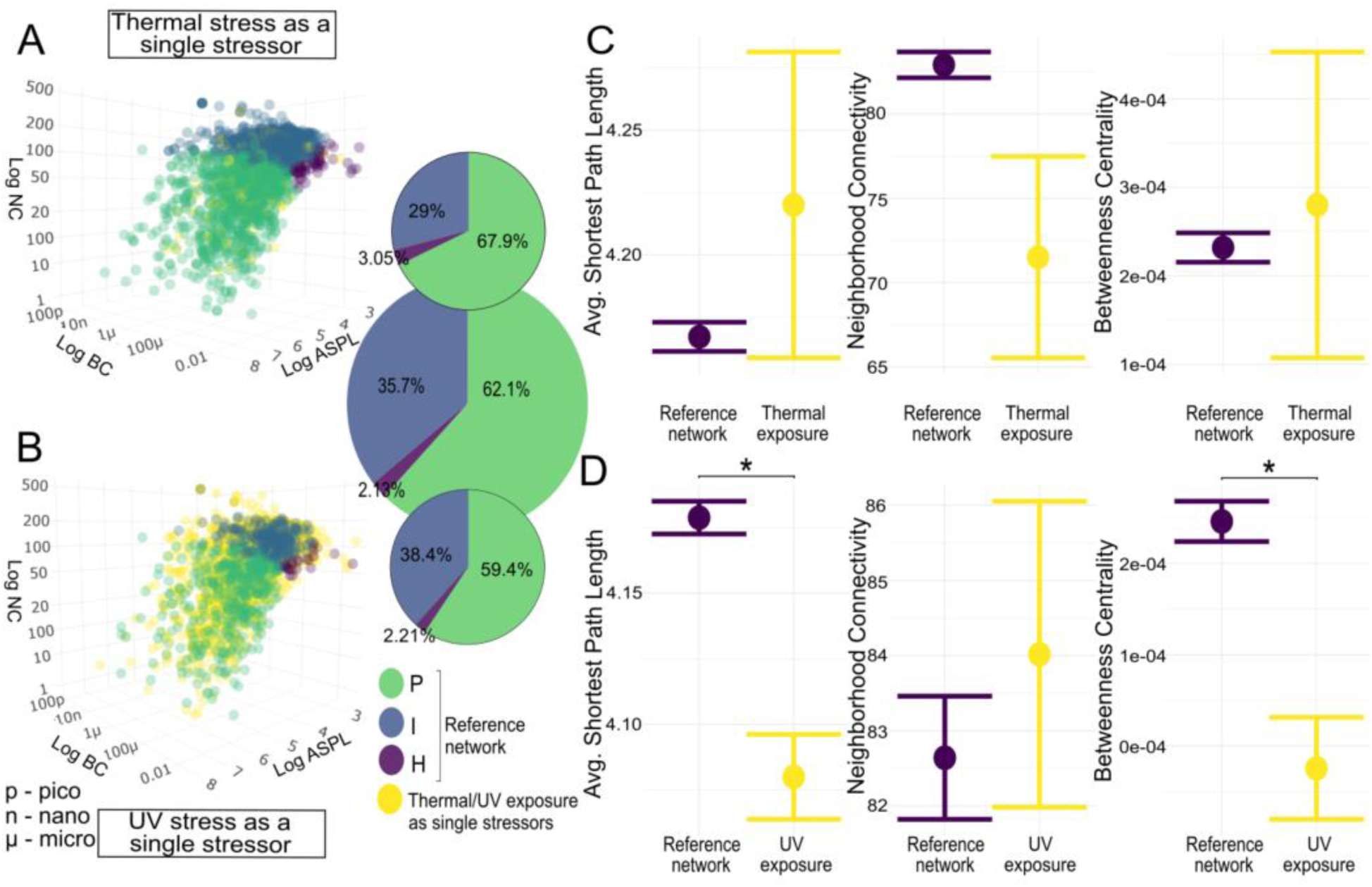
The position of the unique DEG responses to heat and UV treatment comparisons within the zebrafish interactome. (A) Distribution of interactome nodes within network parameter space represented by NC, ASPL, and BC, with node categories shown in green (P nodes), blue (I nodes), and purple (H nodes). (A) TS-unique DEGs and (B) UV-unique DEGs are shown in yellow, respectively. Inset pie charts represent node category distribution (large - interactome; small/top - TS, small/bottom - UV-unique genes). (C, D) Means with standard error plots of ASPL, BC and NC for the interactome and TS and UV - unique gene sets’ network dimensions. Significance levels of pairwise Mann-Whitney U tests are indicated with asterisks p <0.0001 ***; p<0.001 **; p<0.05 *.

### Hypothesis (iv). Differences in DEG positions relate to changes in phenotype between treatment comparisons, and are reflected in fold change

Phenotypic outcomes varied across the three treatment comparisons. In the UV treatment comparison, embryos were generally shorter and had more defects. TS and TS->UV treatment comparisons were similar to one another in terms of (compared to UV) longer embryos with less defects(Fig. 6., Supplementary Data X). The sum of defects and embryo length covaried with ASPL NC, while BC covaried with defect score and embryo length for UV and TS->UV, but not in TS. In TS and TS->UV, defects were low, SEL was high, with high NC and low ASPL. In UV, defects were high, SEL was low, NC high and ASPL low. BC was high in TS, lower in UV, and lowest in TS->UV.

**Fig. 6:**
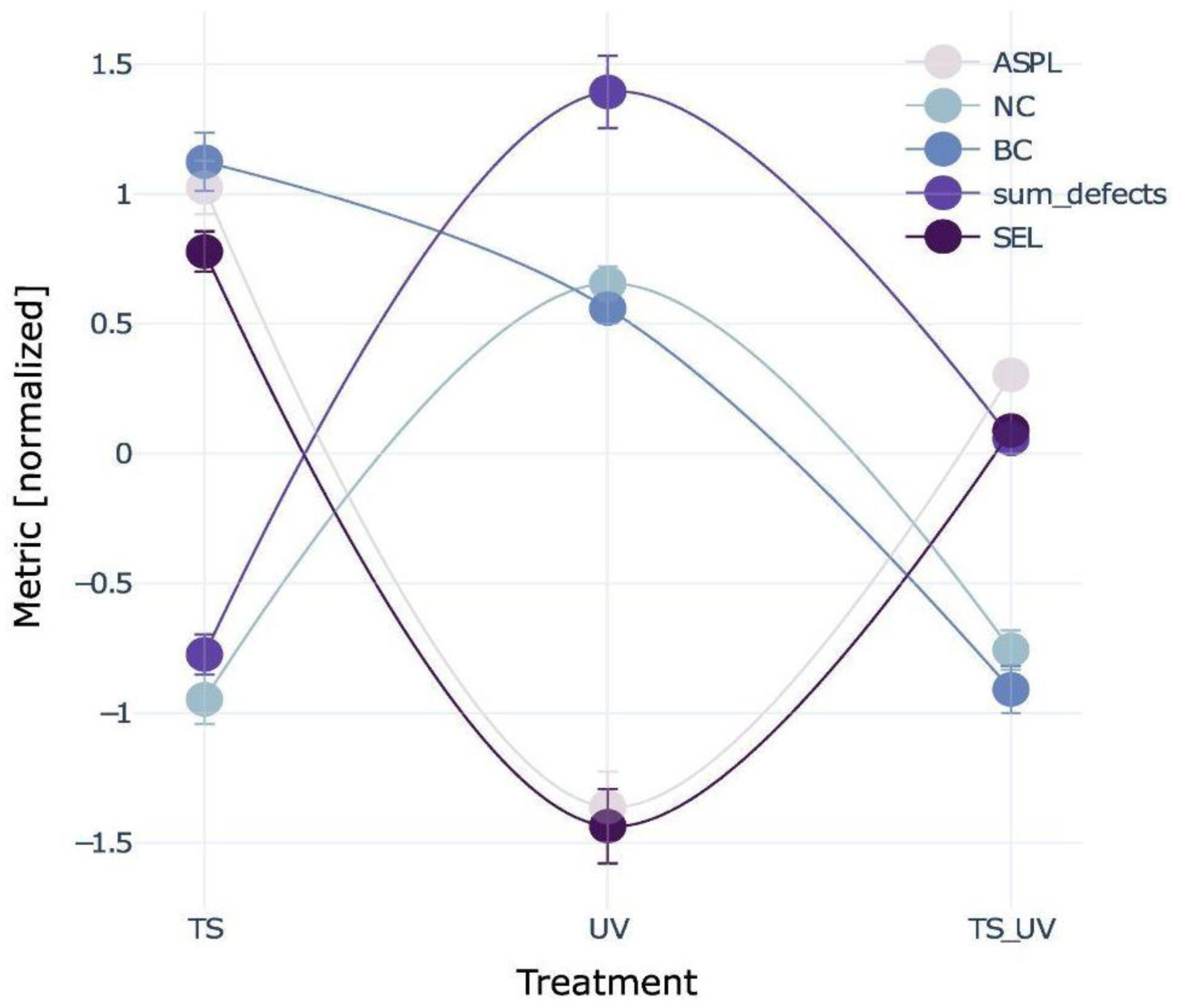
Co-variation of DEG network parameters and phenotypic outcomes. Normalized average values per treatment comparison (with error bars) for interactome-derived network parameters ASPL, NC and BC of DEGs per treatment comparison and phenotypic parameters embryo length and defects. ASPL, NC, sum of defects and SEL co-vary among treatment comparisons while BC shows a different pattern of being highest in TS, lower in UV, and lowest in TS->UV DEGs.

To further explore how phenotypic outcomes may be related to network position of DEGs, we explored significant biological processes, ZFIN phenotypes and ZFIN diseases associated with node categories of DEGs and their commonalities across treatment comparisons (Fig. 7).

**Fig. 7:**
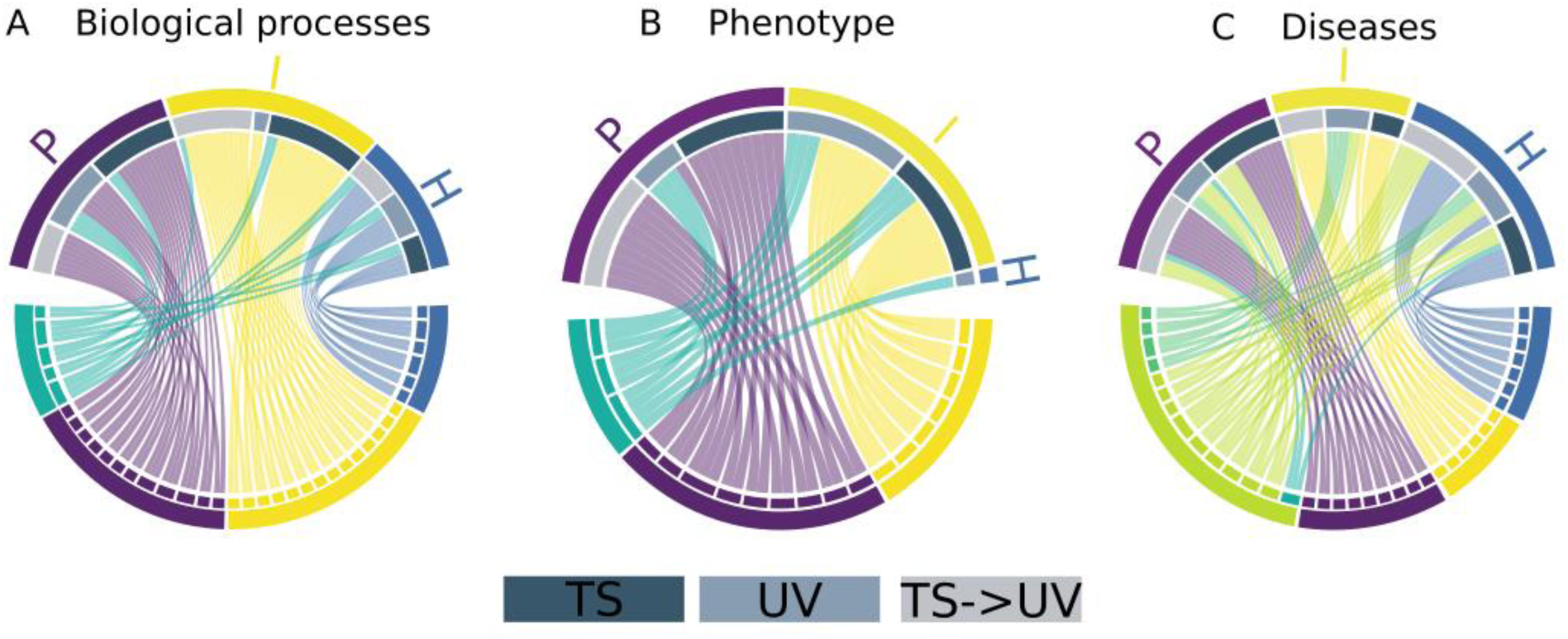
Shared phenotypic effects of DEGs by treatment comparison and network node category. (A -C) Chord diagrams showing shared phenotypic properties of DEGs within the zebrafish interactome (bottom half circle) depending on interactome node location per treatment comparison (top half circle: TS - light gray; UV - medium gray - TS > UV - dark gray). Where (A) shows shared Gene Ontologies (only Biological Process), (B) phenotype and (C) disease implications in zebrafish. Blue ribbons indicate phenotypic properties exclusively associated with H nodes, yellow - I nodes, purple - P nodes. In contrast, Green shades indicate phenotypic properties common across multiple node categories with Turquoise representing all three treatment comparisons; light green representing only two treatment comparison types, and dark green indicating only one treatment comparison type across different node types. Here, For a complete list of significantly enriched terms and genes contributing to them.

Six H-node Biological Processes were common across all treatment comparisons, including carboxylic acid and oxoacid metabolic processes, cellular amino acid biosynthesis, and response to oxidative stress. H nodes of TS and UV treatment comparisons shared family amino acid and small molecule metabolic processes. Fourteen biological processes unique to the I node category included ADP hydrolysis, glycolysis, and nucleoside metabolism shared between TS and TS->UV but not UV, ribonucleoside diphosphate metabolism specific to TS and TS->UV, and ribonucleoprotein and ribosome synthesis between UV and TS->UV. Twelve biological processes in the P node category included three GOs related to cell development and differentiation across all treatments, three GOs on muscle development between TS and TS->UV, and four GOs on eye development and morphogenesis shared between UV and TS->UV. The remaining five biological processes shared between any treatment comparisons were found in multiple node categories. Biological processes, particularly those related to tissue, organ development, and morphogenesis, were shared between H and P nodes across UV and TS->UV treatment comparisons, indicating a cross-category involvement of genes in these specific processes. Processes shared between H and I nodes, including monocarboxylic acid metabolism and carbohydrate catabolism, were associated with UV treatment comparisons, with gene categorizations varying across TS, TS->UV, and UV treatments (Fig. 7).

With regards to phenotype, no phenotype was only associated with DEGs in H nodes. Seven phenotypes related to abnormalities in various body parts, including the central nervous system, gut, liver, and yolk appearance, were uniquely shared between DEGs of I nodes in the UV and TS->UV treatment comparisons. Nine unique phenotypes linked to DEGs in P nodes included disrupted thigmotaxis across all treatment comparisons, and abnormalities in brain volume, spinal structure, and osteoblast accumulation, specifically associated with TS and TS->UV treatments. Three phenotypes - decreased eye size, decreased head size, and edematous pericardium - were common across different treatments but varied in node category associations, such as H nodes in TS, and I and P nodes primarily in UV and TS->UV(Fig. 7, Supplementary Table X).

Seven diseases linked exclusively to the H node category across treatments included diabetic retinopathy associated with TS and UV, while DEGs in H nodes from TS and TS->UV were connected to diseases such as cystitis, melanoma, pancreatic cancer, optic neuritis, pseudoxanthoma elasticum, and vitiligo. There were six diseases associated with only the I node category across multiple treatment comparisons: Sciatic neuropathy was associated with TS and UV. Brain ischemia, congenital nonspherocytic hemolytic anemia, inclusion body myositis, neuromuscular disease and glycogen storage disease were associated with TS and TS->UV (Fig. 7).

There were nine diseases associated with only the P node category across the TS and TS->UV treatment comparisons: heart disease, hereditary spastic paraplegia, nephrogenic diabetes insipidus, oculocutaneous albinism, osteogenesis imperfecta, osteogenesis imperfecta type 4, pigmentation disease, spondylocostal dysostosis, and ullrich congenital muscular dystrophy. Fourteen diseases were associated with multiple node categories across different treatment comparisons. Type 2 diabetes mellitus was unique in being linked to all three treatments—TS and UV in P nodes, and TS->UV in H nodes. Diseases associated with at least two treatment types include cardiomyopathy, cataract, Ehlers-Danlos syndrome, urinary bladder cancer, pulmonary fibrosis, Alzheimer’s disease, asthma, dilated cardiomyopathy, and myocardial infarction. Additionally, diseases associated specifically with the UV treatment and multiple node categories include breast cancer, hypertension, prostate cancer, and renal cell carcinoma (Fig.7).

No common biological processes or phenotypes were observed between the antagonistic and synergistic effects of TS and UV. Diseases linked with the antagonistic DEGs included diabetic retinopathy, associated with P and H nodes, and type 2 diabetes mellitus, associated with P and I nodes. Breast carcinoma was the only disease associated with both effects, with relevant genes found in the H node category.

Fig. 8 shows correlation scatterplots between the squared log fold change of the significant DEGs, and the Biological Process GO terms, ZFIN phenotype components, and ZFIN diseases for each treatment comparison TS, UV and TS->UV. Scatterplots of log fold change of DEGs per treatment comparison reveal two distinct categories: one with low squared log fold change and involvement in a high number of phenotypic effects, and one with a high squared log fold change and involvement in a lower number of phenotypic outcomes (cumulative sum of gene ontologies/biological processes, ZFIN phenotypes or ZFIN diseases per gene). The majority of I and H nodes are in the first group where the latter group is mostly composed of P nodes, with the exception of Phenotype components and phenotype log fold change, which however covers the smallest range of log fold change values among all comparisons. A few H and I genes comprise notable outliers to this pattern, with higher fold changes combined with lower phenotypic involvement; these are *slc25a4* (padj_TS < 0.001, lfc_TS = 2.55, H-node) and *hsc70* (padj_TS = 0.052, lfc_TS= 2.28, I-node) in TS, *hsp70.2* (padj_UV = 2.60E-15, lfc_UV = 6.47, I-node) in UV, and *slc25a4* (padj_TS->UV <0.001, lfc=1.46, I-node) and *grin1b* (padj_TS->UV <0.001, lfc= 1.62, I-node) in TS->UV (Fig. 8).

**Fig. 8:**
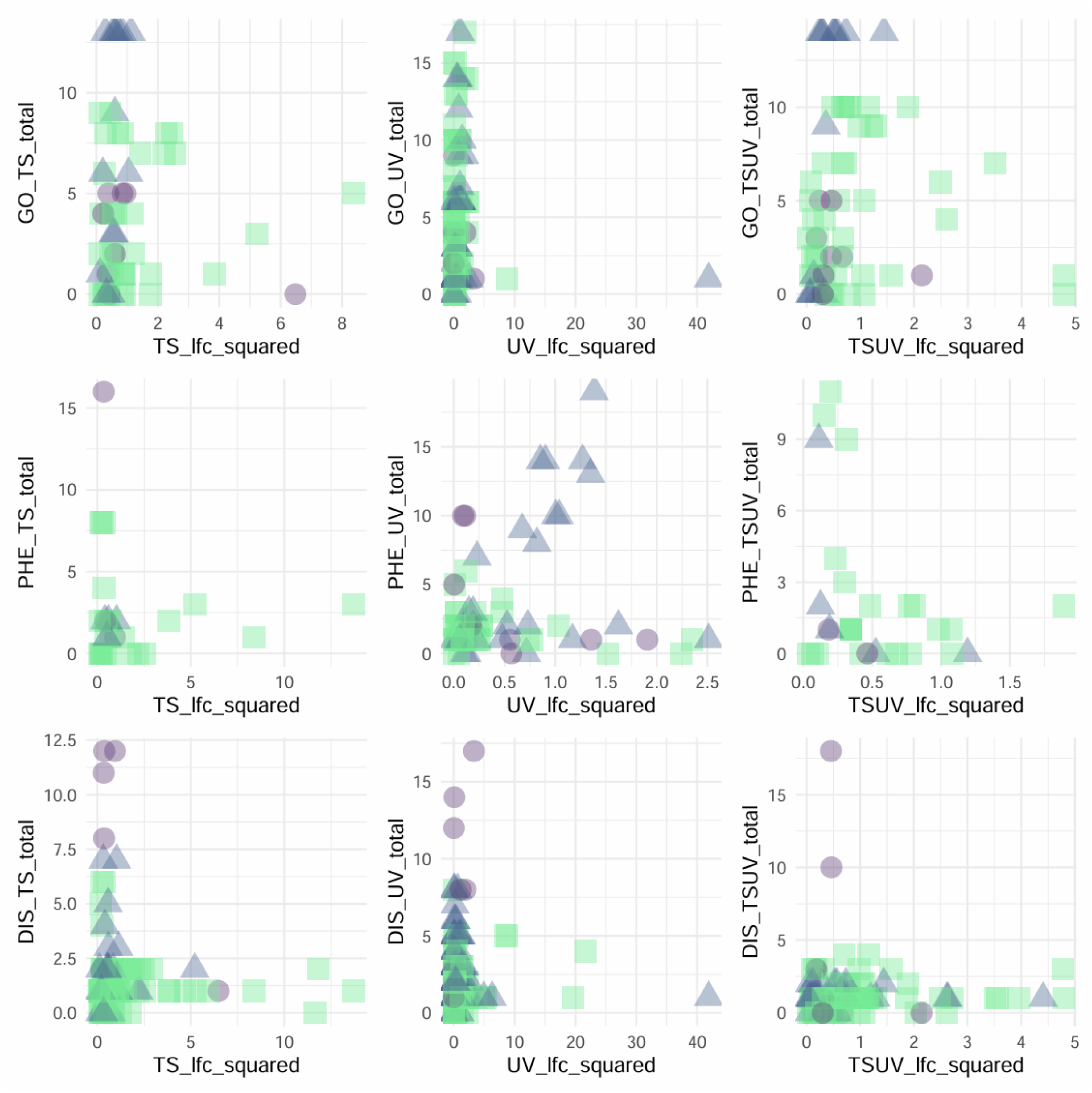
Phenotypic effects of DEGs by treatment comparison and log fold change depending on network node category. Scatter plots of log fold change per treatment comparison with node categories represented with different colors and shapes: I-nodes (blue triangles), H-nodes (purple dots) and P-nodes (green squares).

## Discussion

Biological systems are hierarchical at multiple scales (Schaffer and Ideker 2021), one of which is represented by protein-protein interaction (PPI) networks. With the increase in genomic resources, their possible applications are already broadening for non-model species (Szklarczyk et al. 2023).

The aim of this paper was to provide a model organism analysis of network-related aspects of the transcriptomic response to environmental stressors, using heat and UV radiation in zebrafish embryos.

### Stress response genes are located centrally in the zebrafish interactome

Genes with stress-related ontologies were significantly more prevalent in both the intermediate and central nodes of the interactome, providing support for **hypothesis (i)**. This central localization confirms both high evolutionary constraint and the potential for robust gene expression under stress, as found in previous studies (Romero and Gormally 2019; Costa-Mattioli and Walter 2020). Even though the general stress response should be highly conserved, the activation and functional consequences of it in specific contexts may be more variable (Romero and Gormally 2019). In our experiment with heat and UVR, 71 genes were consistently expressed across all treatments *versus* their respective controls, supporting the results from **hypothesis (i)** of a core set of stress-responsive genes that may mediate a generalized response *via* multiple pathways across the network (de Nadal, Ammerer, and Posas 2011). However, the relative positions of DEG groups *per* treatment within the zebrafish interactome were not consistently observed to be more central as expected under **hypothesis (ii)**. While all three treatments activated a higher number of DEGs in hub nodes, thus reflecting a central component, many DEGs in response to UV exposure were located in intermediate nodes. Heat preceding UV treatment overall induced both more hub and intermediate genes, while both TS and Heat preceding UV treatments both increased DEGs in hub and peripheral nodes compared to UV, indicating a treatment-specific response pattern.

The higher number of peripheral nodes compared to intermediate and hub nodes in both treatments involving heat stress suggests that, while the general stress response may be conserved, multiple components of the network may be activated in response to different external stressors (McClintock 1984). This perhaps reflects a trade-off between evolutionary robustness constraint and stressor-specific gene expression, an architectural requirement of evolvable systems (Kitano 2004) and a trait also seen in computational simulations of evolved artificial gene regulatory networks (Turner et al. 2019; Turner and Wollenberg Valero 2021)

The intermediate position of nodes responding to UV indicates that the innate response to UV is regulated *via* nodes that affect many other gene products through having the highest neighborhood connectivity (Wollenberg Valero 2020), which may cause activation of multiple genes in a pathway, potentially leading to survival consequences (Jeong et al. 2001; Bonatto 2007). It needs to be explored further whether the differentially expressed genes are related to repair or damage processes, or both (Zagarese and Williamson 2001; Dong et al. 2007; Chen et al. 2020; Feugere et al., n.d.).

Interestingly, the genes expressed by zebrafish embryos in this experiment were not a representative subset of the network topology of the overall zebrafish interactome, containing significantly fewer hub and peripheral nodes and substantially more intermediate nodes. This pattern indicates that development may predominantly be regulated by intermediately located genes that have many functional connections with each other (Wollenberg Valero 2020). However, even within this fundamentally different node distribution, the effects of the three treatment comparisons were the same as when compared to the zebrafish interactome: the UV exposure was mostly characterized by DEGs in intermediate and hub nodes, whereas any addition of heat either in TS or in TS->UV led to activation of less intermediate and more peripheral nodes. This underscores the idea that the response to thermal stress activates both central and peripheral portions of the network.

### Uniform network positions of genes activated by different stressor combinations

We found distinct expression patterns under different stress conditions. UV exposure (in the form of a UVB/UVA damage/repair assay, (Dong et al. 2007)) resulted in the highest number of expressed genes, indicating a strong genetic response to UV challenge. Comparatively, thermal stress followed by UV exposure (TS->UV) compared to UV exposure without thermal stress had the second highest number of differentially expressed genes, many of which were different from either the UV or TS treatments by themselves. This suggests an non-linear effect of these sequential stress exposures on gene expression. However, network topological differences were not significant in genes synergistically active in TS->UV relative to the interactome background. Some genes only responding synergistically in TS->UV were located in the network periphery, which do not support the hypothesis that synergistic genes would be more central due to the need to mount a general stress response. There likewise was no significant difference in the position of antagonistic DEGs compared to the interactome, but eight of these genes were located in the periphery. These genes may represent stressor-unique effects being located in peripheral locations not being differently activated when stressors are combined, but due to missing statistical significance does not support hypothesis 3, either. Looking at stressor-specific genes, TS-specific DEGs did not differ statistically from the overall distribution of nodes in the interactome, whereas DEGs unique to UVR did differ significantly in both average shortest path length and neighborhood connectivity. UVR activated five more intermediate nodes, which overall does not conclusively support the hypothesis 3 that unique responses to single stressors are more peripheral. In addition, no common biological processes or phenotypes were found between the antagonistic and synergistic effects of heat and UVR, except for breast carcinoma, which was associated with both antagonistic and synergistic gene sets linked to the hub node category, while diabetic retinopathy and type 2 diabetes mellitus were linked to different node categories in the antagonistic set of DEGs.

### DEG positions are linked to phenotypic effects and expression fold changes

In our experiment, pre-exposing embryos to heat before the UVR damage/repair assay led to fewer defects compared to direct UV exposure alone. We found a pattern of covariation between the number of embryo defects and specific network parameters of DEGs: UV treatment, which caused the highest number of defects, was associated with DEGs that had a relatively low average shortest path length and high neighborhood connectivity, reflecting the increase in intermediate nodes described above. In contrast, the heat stress (TS) and heat followed by UV exposure (TS->UV) treatments showed the opposite trends. These correlative findings suggest that the network position of DEGs related to each treatment may influence phenotypic outcomes, with the addition of heat causing a shift to DEGs more peripherally in the network, while lowering negative phenotypic outcomes compared to UV only.

Besides directly measuring phenotypic outcomes, different stress treatments activated several genes related to identical biological processes, phenotypes and diseases, in each node category. This indicates that besides a central, conserved portion of the stress response (Romero and Gormally 2019), also parts of the network associated with intermediate and peripheral nodes can contribute to stress response independently of the specific stressor. Our results provide evidence for an evolutionary conserved component of the stress response in terms of organismal development and function, which can limit the risk of deleterious effects in mismatched environments (de Nadal, Ammerer, and Posas 2011; Burton et al. 2021). This conservation exists across multiple node categories of the network, and adaptability may be built-in at the periphery of the network, which allows different genes to be modified in response to different stressors, but still affecting the same phenotype components. Heat and UV have previously been shown to activate many of the same pathways as when exposed to either separate treatment in plants (Jenkins, Suzuki, and Mount 1997; Jardim et al. 2015).

Specifically, across all three treatments, hub nodes were linked to oxidative stress response and amino acid biosynthesis. These genes are known to involve a variety of phenotypic abnormalities including cancers, suggesting a critical role in managing cellular stress and metabolic processes. The general stress response is activated under heat stress in zebrafish embryos (Long et al. 2012). Both heat and UV can induce oxidative stress in cells, as a result of which protein synthesis is lowered via the eIF2α and ATF4 pathways (Harding et al. 2003). Intermediate nodes of all treatments were related to energy management and the synthesis of DNA/RNA and proteins under stress, while diseases associated with DEGs in intermediate nodes also affect cellular damage and stress, both in muscles and nerves. These processes are aligned to the energetic costs of the stress response (Picard et al. 2018).

Peripheral nodes across treatments were linked to structural and morphological development, indicating their role in anatomical and developmental pathways. DEGs located in peripheral nodes across all treatments affected the response to touch, and the formation of a straight spine with proper volumes of blood and CNS fluids. These observations relate to the idea that peripheral nodes, which show the least amount of pleiotropic interactions, are involved in stressor-specific outcomes in defined components of the phenotype (Wollenberg Valero 2020). Spine malformation “Bent spine” is a common developmental malformation in zebrafish (von Hellfeld et al. 2020) and has, along with an altered behavioral response to stimuli, has been observed in response to heat and UV treatment in zebrafish embryos and larvae (Feugere et al. 2021, 2023, n.d.).

One observed stressor-specific response was the higher proportion of peripheral node DEGs across the two treatments involving heat (TS and TS->UV), with a number of genetic disorders affecting a wide range of organ systems, indicating that thermal stress may systemically affect many structural and developmental genetic pathways. Multiple systems are known to be activated in zebrafish by thermal stress such as the innate immune response (Zhang et al. 2018), reactive oxygen species accumulation (Luo et al. 2015; Mariana and Badr 2019), and a generally faster developmental speed (Feugere et al., n.d.). Heat activates a number of regulatory processes such as nucleotide and ribonucleotide metabolic processes which are related to the general small molecule metabolic activity of the cell. This is metabolic priming is a possible mechanism for thermal stress prior to UV-exposure resulting in a cross-protective, hormetic effect, where for example repair mechanisms or other transcription processes are upregulated (Costantini, Metcalfe, and Monaghan 2010; Rattan 2006; Feugere et al., n.d.; Rodgers and Gomez Isaza 2023) Evidence for a hormetic effect is reflected in the phenotypic outcomes, specifically the occurrence of defects which was lowered in the thermal stress followed by UV treatment comparison compared to UV alone.

Genes related to morphological structure formation and involved in response to UV were located both in peripheral and hub nodes which may represent specific developmental pathways that are regulated centrally, but with specific phenotypic effects (Singh et al. 2008). Diseases associated with the UV treatment across multiple node categories involve cell growth and division and the cardiovascular system, which underscores the nucleic acid damage and resulting impacts on the cell cycle inflicted by UVR (Bootsma and Humphrey 1968). In both UV and TS->UV treatments, differentially expressed genes with consequences for early embryonic development were associated with intermediate nodes. Across all node categories, DEGs in these treatments further shared decreased head size and pericardial edema phenotypes, which are typical for UV radiation-induced damages in zebrafish (Alves and Agustí 2020).

In addition to the position of stress-induced DEGs in the interactome, we looked at the interplay between node position, phenotype involvement, and gene expression fold changes. DEGs had either low log fold changes and were associated with numerous phenotypic effects, or had high log fold changes and were linked to fewer phenotypic outcomes. Genes in the first category represented most intermediate and hub nodes, while peripheral node genes comprised most genes in the second category. Genes with large phenotypic effects are known to be pleiotropic, which explains why H and I nodes were associated with multiple phenotypes (Wang, Liao, and Zhang 2010; Fraser et al. 2002). This pattern has previously been dubbed the “cost of complexity” (Wagner et al. 2008; Orr 2000). The “E-R anticorrelation” expects (central H and pleiotropic I-nodes) genes that are highly constrained to be highly expressed (Joy et al. 2005; Wollenberg Valero 2020). Fold changes in such highly expressed genes may have effects on more essential components of the phenotype, even if they are lower in magnitude. Conversely, genes located peripherally may tolerate larger magnitude of expression changes to show distinct phenotypic effects which could explain this observed pattern. Constraint imposed by protein-protein interactions has previously been found to influence protein expression responses to rearing temperature in fish adapted to warm and cold streams, with high numbers of protein-protein interactions lowering protein expression (Papakostas et al. 2014).

Nonetheless, we found a few intermediate and hub genes to constitute outliers to this pattern, showing higher expression changes combined with lower phenotypic involvement. These were the two heat shock protein genes *hsc70* (synonymous with *hspa8*; heat shock cognate 70, in TS) and *hsp70.2* in UV. *hsc70* is a constitutive molecular chaperone maintaining protein homeostasis, active in development (Santacruz, Vriz, and Angelier 1997; Stelzer et al. 2016), but is known to be upregulated in response to heavy metal exposure in developing sea urchin embryos (Pinsino et al. 2011), and in heat stress-induced transposable element activation in the germline (Cappucci et al. 2019). *Hsp70.2* is orthologous to human heat shock protein 70, with molecular chaperone function and is inducible by heat (Lele, Engel, and Krone 1997; Stelzer et al. 2016). In treatments with heat stress (TS and TS->UV), *slc25a4* (Solute Carrier Family 25 Member 4) constituted another such outlier. This antiporter translocates ADP into the mitochondrial matrix and ATP back into the cytoplasm (Stelzer et al. 2016; Thompson et al. 2016), and so could play a major role in energy availability for the stress response (Picard et al. 2018). Lastly *grin1b* in TS->UV (Glutamate Ionotropic Receptor NMDA Type Subunit 1) codes for a glutamate receptor which is involved in synapse plasticity, but also MAPK signaling (Stelzer et al. 2016) and is known to affect behavior in developing zebrafish (Zoodsma et al. 2020). Overall, the roles of *slc25a4* and *grin1b* in the stress response are little explored, compared to the two heat shock proteins. Comparing fold changes to network position may therefore aid in identifying key regulators of the cellular and physiological systemic response to various stressors.

## Conclusions

Overall our results support the idea of the response to different environmental stressors being evolutionarily conserved and central to the zebrafish interactome, but also including some peripheral parts of the network. hub, intermediate and peripheral network positions of DEGs represented different aspects of the stress response. There were no similarities between DEG sets that responded to individual stressors. Combining network position and fold changes in response to treatment may aid in identifying novel regulators of the response to specific stressors. DEGs could clearly be linked to developmental outcomes, evident in phenotypes like bent spine, decreased eye size, and pericardial edema. Since the embryonic environment can enhance, impede, or generally change later-life phenotypic plasticity, impacting the individual’s ability to acclimate to environmental stressors (Scott and Johnston 2012; Beaman, White, and Seebacher 2016; Loughland, Little, and Seebacher 2021), research on stress responses in aquatic developing organisms should receive increased attention in the context of climate change affecting aquatic environments.

## Acknowledgments

This work was funded by the Royal Society (RGS\R2\180033) to KWV. KWV and KB acknowledge funding by the European Union (ERC, MolStressH2O, 101044202). Views and opinions expressed are however those of the author(s) only and do not necessarily reflect those of the European Union or the European Research Council Executive Agency. Neither the European Union nor the granting authority can be held responsible for them. Furthermore, we acknowledge the Viper High Performance Computing facility of the University of Hull and its support team.

## Supplementary Materials to the manuscript

**Supplementary Fig S1:**
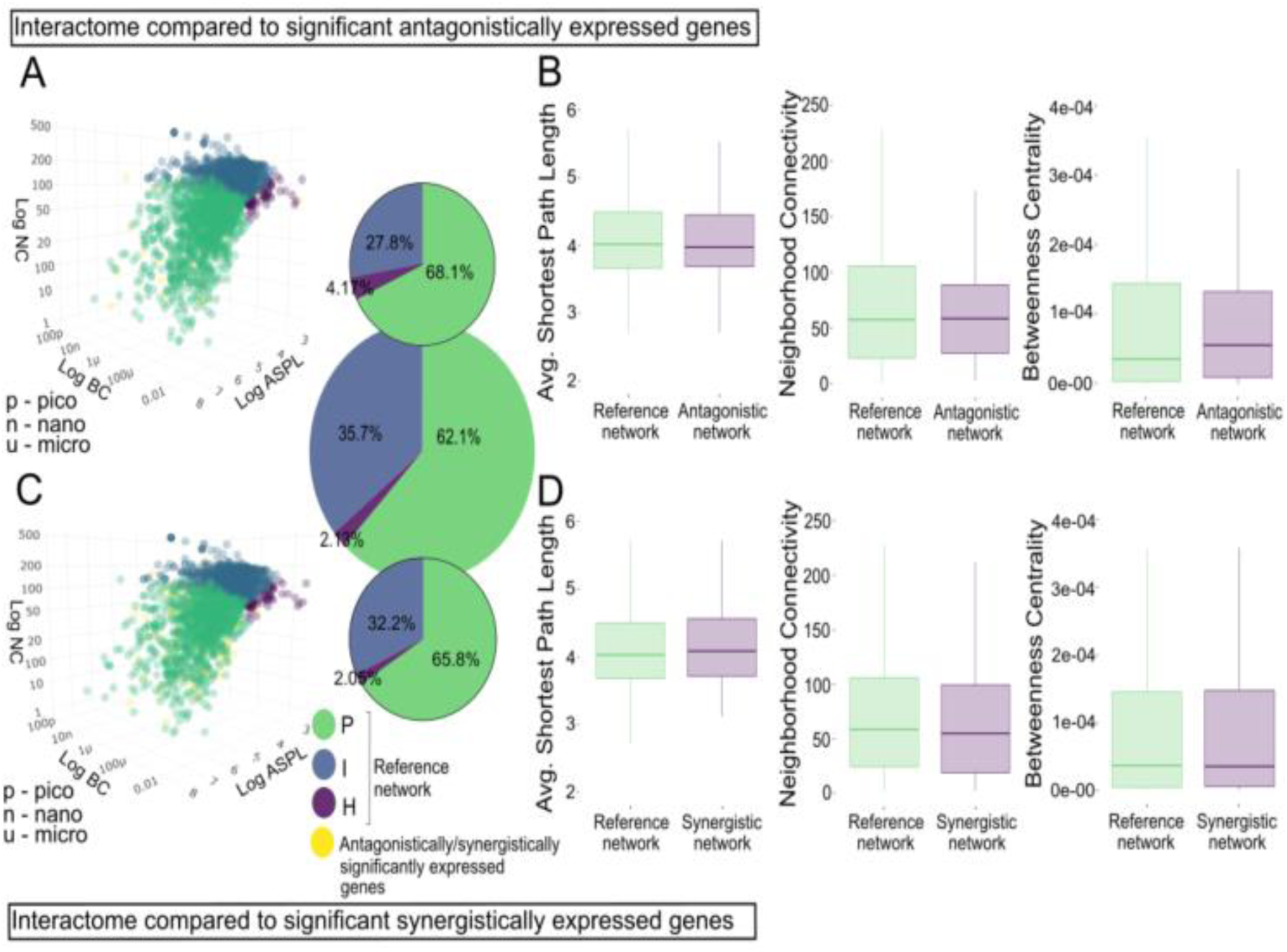
The position of synergistic and antagonistic DEGs within the zebrafish interactome. (A) Distribution of interactome network nodes within network parameter space represented by NC, ASPL, and BC, with node categories shown in green (P nodes), blue (I nodes), and purple (H nodes). (A) Antagonistic genes and (C) Synergistic genes are shown in yellow, respectively. Inset pie charts represent node category distribution (large - interactome; small/top - antagonistic, small/bottom - synergistic genes). (B,D) Error plos of ASPL, BC and NC for the interactome and synergistic and antagonistic gene sets’ network dimensions. Pairwise Mann-Whitney U tests were not significant.

**Supplementary Table S1.** Independent-sample Kruskal-Wallis test summary for differences between node types (interactome vs. genes with “stress response” GO term) for the three network statistical parameters. The test statistic is adjusted for ties. ASPL - Average Shortest Path Length; NC - Neighborhood Connectivity; BC - Betweenness Centrality. Corresponding with Fig. 1B.

| Network statistical parameter | Total N | Test Statistic KW-H | Degree of Freedom | Asymptotic Sig. (2-sided test) | eta2[H] |
| --- | --- | --- | --- | --- | --- |
| ASPL | 13791 | 122.454 | 1 | <0.001 | 0.008 |
| NC | 13791 | 0.000 | 1 | 0.988 | -0.00007 |
| BC | 11187 | 127.626 | 1 | <0.001 | 0.011 |

**Supplementary Table S2.** Independent-sample Kruskal-Wallis test summary for differences between node types (interactome vs. experimental DEGs) for the three network statistical parameters. The test statistic is adjusted for ties. ASPL - Average Shortest Path Length; NC - Neighborhood Connectivity; BC - Betweenness Centrality. Corresponding with Fig. 3 (A-C).

| Network statistical parameter | Total N | Test Statistic KW-H | Degree of Freedom | Asymptotic Sig. (2-sided test) | eta2[H] |
| --- | --- | --- | --- | --- | --- |
| ASPL | 14360 | 20.778 | 3 | <0.001 | 0.001 |
| NC | 14360 | 19.837 | 3 | <0.001 | 0.001 |
| BC | 11658 | 19.795 | 3 | <0.001 | 0.001 |

**Supplementary Table S2.1.** ASPL Pairwise comparisons from Mann-Whitney Wilcoxon tests. Tests for difference in average shortest path length between DEGs from the different treatment comparisons against the interactome. All significance values have been adjusted by Bonferroni correction for multiple tests. Corresponding with Fig. 3A.

| Gene set 1 | Gene set 2 | W | Std. Error | Std. W | P | Padj. |
| --- | --- | --- | --- | --- | --- | --- |
| interactome | TS | -217.451 | 199.739 | -1.089 | 0.276 | 1 |
| interactome | UV | 420.675 | 99.165 | 4.242 | 0 | 0 |
| interactome | TS->UV | -88.624 | 167.256 | -0.53 | 0.596 | 1 |

**Supplementary Table S2.2.** NC Pairwise comparisons from Mann-Whitney Wilcoxon tests. Tests for difference in neighborhood connectivity between DEGs from the different treatment comparisons against the interactome. All significance values have been adjusted by Bonferroni correction for multiple tests. Corresponding with Fig. 3B.

| Gene set 1 | Gene set 2 | W | Std. Error | Std. W | P | Padj. |
| --- | --- | --- | --- | --- | --- | --- |
| interactome | TS | 630.619 | 199.739 | 3.157 | 0.002 | 0.01 |
| interactome | UV | -130.705 | 99.165 | -1.318 | 0.187 | 1 |
| interactome | TS->UV | 453.921 | 167.256 | 2.714 | 0.007 | 0.04 |

**Supplementary Table S2.3.** BC Pairwise comparisons from Mann-Whitney Wilcoxon tests. Tests for difference in betweenness centrality between DEGs from the different treatment comparisons against the interactome. All significance values have been adjusted by Bonferroni correction for multiple tests. Corresponding with Fig. 3C.

| Gene set 1 | Gene set 2 | W | Std. Error | Std. W | P | Padj. |
| --- | --- | --- | --- | --- | --- | --- |
| interactome | TS | -525.452 | 178.755 | -2.94 | 0.003 | 0.02 |
| interactome | UV | -158.669 | 87.9 | -1.805 | 0.071 | 0.426 |
| interactome | TS->UV | -481.9 | 150.028 | -3.212 | 0.001 | 0.008 |

**Supplementary Table S3.** Independent-sample Kruskal-Wallis test summary for differences between node types (expressed network vs. experimental DEGs) for the three network statistical parameters. The test statistic is adjusted for ties. ASPL - Average Shortest Path Length; NC - Neighborhood Connectivity; BC - Betweenness Centrality. Corresponding with Fig 4 (A-C).

| Network statistical parameter | Total N | Test Statistic KW-H | Degree of Freedom | Asymptotic Sig. (2-sided test) | eta2[H] |
| --- | --- | --- | --- | --- | --- |
| ASPL | 11321 | 13.226 | 3 | 0.004 | 0.0009 |
| NC | 11321 | 20.253 | 3 | <0.001 | 0.002 |
| BC | 9395 | 6.297 | 3 | 0.098 | 0.0004 |

**Supplementary Table S3.1.** ASPL Pairwise comparisons from Mann-Whitney Wilcoxon tests. Tests for difference in average shortest path length between DEGs from the different treatment comparisons against the expressed network. Corresponding with Fig. 4A.

| Gene set 1 | Gene set 2 | W | Std. Error | Std. W | P | Padj. |
| --- | --- | --- | --- | --- | --- | --- |
| Expressed network | TS | -341.475 | 158.596 | -2.153 | 0.031 | 0.188 |
| Expressed network | UV | 154.906 | 80.423 | 1.926 | 0.054 | 0.325 |
| Expressed network | TS->UV | -246.017 | 133.203 | -1.847 | 0.065 | 0.389 |

**Supplementary Table S3.2.**
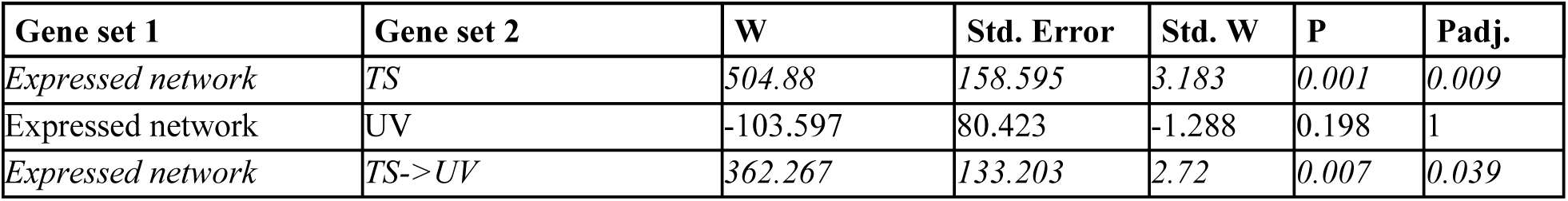
NC Pairwise comparisons from Mann-Whitney Wilcoxon tests. Tests for difference in neighborhood connectivity between DEGs from the different treatment comparisons against the expressed network. Corresponding with Fig. 4B.

| Gene set 1 | Gene set 2 | W | Std. Error | Std. W | P | Padj. |
| --- | --- | --- | --- | --- | --- | --- |
| Expressed network | TS | 504.88 | 158.595 | 3.183 | 0.001 | 0.009 |
| Expressed network | UV | -103.597 | 80.423 | -1.288 | 0.198 | 1 |
| Expressed network | TS->UV | 362.267 | 133.203 | 2.72 | 0.007 | 0.039 |

**Supplementary Table S4.** Antagonistic and synergistic independent-sample Kruskal-Wallis test summary for differences between node types (interactome vs. antagonistic/synergistic) for the three statistical parameters. The test statistic is adjusted for ties. ASPL - Average Shortest Path Length; NC - Neighborhood Connectivity; BC - Betweenness Centrality. The significance level is .050. Corresponding with Fig. 6 (B and D).

| Network statistical parameter | Total N | Test Statistic KW-H | Degree of Freedom | Asymptotic Sig. (2-sided test) | eta2[H] |
| --- | --- | --- | --- | --- | --- |
| **Antagonistic** |  |  |  |  |  |
| ASPL | 13791 | .000 | 1 | .991 | -7.25E-05 |
| NC | 13791 | 1.340 | 1 | .247 | 2.47E-05 |
| BC | 13791 | .010 | 1 | .922 | -7.18E-05 |
| **Synergistic** |  |  |  |  |  |
| ASPL | 13791 | 1.235 | 1 | .266 | 1.70E-05 |
| NC | 13791 | 3.129 | 1 | .077 | 0.0002 |
| BC | 13791 | .297 | 1 | .586 | -5.10E-05 |

**Supplementary Table S5.**
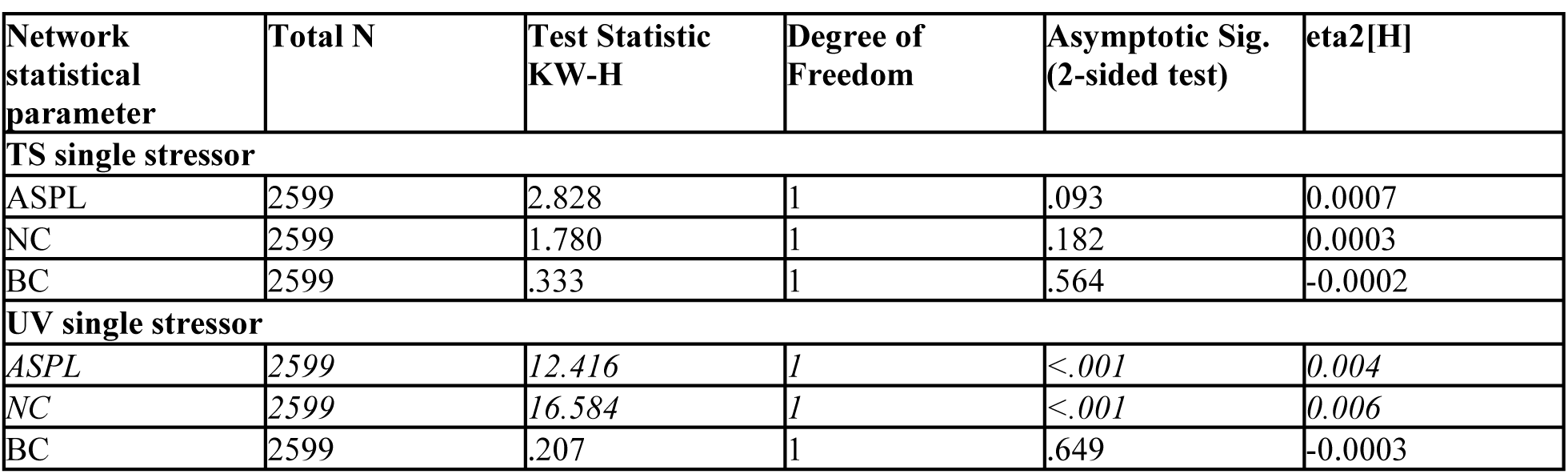
Thermal stress and UV exposure as single stressors independent-sample Kruskal-Wallis test summary for differences between node types (interactome vs. single stressors) for the three statistical parameters. The test statistic is adjusted for ties. ASPL - Average Shortest Path Length; NC - Neighborhood Connectivity; BC - Betweenness Centrality. The significance level is .050. Corresponding with Fig. 6 (B and D).

| Network statistical parameter | Total N | Test Statistic KW-H | Degree of Freedom | Asymptotic Sig. (2-sided test) | eta2[H] |
| --- | --- | --- | --- | --- | --- |
| **TS single stressor** |  |  |  |  |  |
| ASPL | 2599 | 2.828 | 1 | .093 | 0.0007 |
| NC | 2599 | 1.780 | 1 | .182 | 0.0003 |
| BC | 2599 | .333 | 1 | .564 | -0.0002 |
| **UV single stressor** |  |  |  |  |  |
| ASPL | 2599 | 12.416 | 1 | <.001 | 0.004 |
| NC | 2599 | 16.584 | 1 | <.001 | 0.006 |
| BC | 2599 | .207 | 1 | .649 | -0.0003 |

**Supplementary Table S6:** Pie charts in Fig 2-6. Table showing node associations between networks.

| Comparison | Figure # | Experimental comparison group | Percentage of targeted H nodes, Total H node percentage, Difference in number of nodes | Percentage of targeted I nodes, Total I node percentage, Difference in number of nodes | Percentage of targeted P nodes, Total P node percentage, Difference in number of nodes |
| --- | --- | --- | --- | --- | --- |
| Interactome | Fig. 1<br>Fig. 3<br>Fig. 5<br>Fig. S1 |  | 2.13%, | 35.7%, | 62.1%, |
| stress GO term genes vs. interactome | Fig. 1 | Stress response (GO term)<br>N= 1176 | 4.19%,<br>+2.06%,<br>+24 | 37%,<br>+1.3%,<br>+15 | 58.9%,<br>-3.2%,<br>-38 |
| Experimental DEGs vs interactome | Fig. 3 | Thermal stress<br>N=461 | 3.79%,<br>+1.33%,<br>+6 | 29.7%,<br>-6%,<br>-27 | 66.5%,<br>+4.4%,<br>+20 |
|  |  | UV exposure<br>N=2113 | 2.41%,<br>+0.28%,<br>+6 | 37.6%,<br>+1.9%,<br>+40 | 60%,<br>-2.1%,<br>-44 |
|  |  | UV preceded by<br>thermal stress<br>N=666 | 2.46%,<br>+0.33%,<br>+2 | 33.2%,<br>-2.5%,<br>-17 | 64.3%,<br>+2.2%,<br>+15 |
| **Expressed network** | **Fig. 4** |  | 0.25% | 74% | 25.7% |
|  |  | Expressed network vs<br>interactome | -1.88%, | +38.3%, | -36.4%, |
| **Experimental DEGs vs.**<br>**expressed network** | **Fig. 4** | Thermal stress<br>N=461 | 0.445%,<br>+0.2%,<br>+ 1 | 66.8%,<br>-7.2%,<br>-33 | 32.7%,<br>+7.07%,<br>+33 |
|  |  | UV exposure<br>N=2113 | 0.386%,<br>+0.059%,<br>+1 | 75%,<br>+1%,<br>+21 | 24.6%,<br>-1.1%,<br>-23 |
|  |  | UV preceded by<br>thermal stress<br>N=666 | 0%,<br>-0.25%,<br>-2 | 70.7%,<br>-3.3%,<br>-22 | 29.3%,<br>+3.6%,<br>+24 |
| **Stressor-unique DEGs**<br>**and interactome** | **Fig. 5** | TS single stressor<br>N=134 | 3.05%,<br>+0.92%,<br>+1 | 29%,<br>-6.7%,<br>-9 | 67.9%,<br>+5.8%,<br>+8 |
|  |  | UV single stressor<br>N=1720 | 2.21%,<br>-0.08%,<br>0 | 38.4%,<br>+2.7%,<br>+5 | 59.4%,<br>-2.7%,<br>-5 |
| **Antagonistic and**<br>**synergistic genes vs**<br>**interactome** | **Fig. S1** | Antagonistic<br>N=135 | 4.17%,<br>+2.04%,<br>+2.8 | 27.8%,<br>-7.9%,<br>-11 | 68.1%,<br>+6%,<br>+8 |
|  |  | Synergistic<br>N=287 | 2.05%,<br>-0.08%,<br>0 | 32.2%,<br>-3.5%,<br>-10 | 65.8%,<br>+3.7%,<br>+11 |

